# Simultaneously cholinergic projection in Ascending and Descending Circuits from Midbrain

**DOI:** 10.1101/2022.06.05.494860

**Authors:** Peilin Zhao, Tao Jiang, Huading Wang, Xueyan Jia, Anan Li, Jing Yuan, Qingming Luo, Xiangning Li, Hui Gong

## Abstract

The midbrain participates in complex neural information processing in the ascending and descending circuits, but their organization remains unclear due to the lack of comprehensive dissection of the characterization of individual neurons. Combining fluorescent micro-optical sectional tomography with sparse labeling, we acquired the whole-brain dataset with high resolution and reconstructed the detailed morphology of the pontine-tegmental cholinergic neurons (PTCNs). As the main cholinergic system of the midbrain, the individual PTCNs own abundant axons with length up to 60 cm and 5000 terminal branches and innervate multiple brain regions from the spinal cord to cortex in both hemispheres. According to various targeting regions in the ascending and descending circuits, individual PTCNs could be grouped into four types and the axonal fibers of cholinergic neurons in the pedunculopontine nucleus present more divergent while neurons in the laterodorsal tegmental nucleus contain richer axonal fibers and dendrites. In the axonal targeting nuclei, such as in the thalamus or cortex, the individual neurons innervate multiple sub-regions with separate pathways. These results provide the detailed organization characterization of the cholinergic neurons to understand the connection logic of the midbrain.

## Introduction

The central nervous system is a highly ordered structure, in which the external information is transmitted to the higher center through the ascending pathway, and the decision-making information is delivered through the descending pathway to determine the physiological changes and behavioral responses of the individuals (L. Luo, 2015). The axonal fibers determine where the neurons transmitted information to while their connection patterns decide the role in the circuit from upstream to downstream regions. The midbrain locates between the forebrain and hindbrain with dense connections involve in ascending and descending circuits among different regions from the spinal cord to the cortex (“The Organization of the Central Nervous System,” 2014). Previous studies have indicated that some specialized populations of midbrain neurons preferentially innervate ascending or descending circuits respectively. DA neurons in the ventral tegmental area (VTA) (Beier et al., 2015) and different types of neurons in the dorsal raphe (DR) (Xu et al., 2021) mainly send their fibers to ascending areas, while the mesencephalic locomotor region modulates the ascending and descending circuits to participate in different functions via two separate group glutamatergic neurons (Ferreira-Pinto et al., 2021). However, cholinergic neurons in the midbrain send abundant axonal projections both in ascending and descending circuits (Henrich Martin et al., 2020; Peilin Zhao et al., 2022), and their projection patterns have not been systematically characterized until now.

The midbrain cholinergic neurons mainly gather in the pedunculopontine nucleus (PPN) and laterodorsal tegmental nucleus (LDT), the two subareas of the pontine-tegmental cholinergic system with different projection patterns (Henrich Martin et al., 2020; I. Huerta-Ocampo et al., 2020; Mena-Segovia et al., 2017; M. Mesulam et al., 1983). As the major source of acetylcholine in many subcortical nuclei, the pontine-tegmental cholinergic neurons (PTCNs) send abundant fibers in three major trajectories (Mena-Segovia & Bolam, 2017) and involve in various functions (Vitale et al., 2019). In the ascending dorsal circuits, PTCNs mediate prefrontal serotonin releasing from the DR (Kinoshita et al., 2018) and participate in multiple functions including auditory sensation (F. Luo et al., 2013), sensorimotor (Muller et al., 2013), and spatial memory (Mitchell et al., 2002) via targeting different thalamic nuclei. In addition, PTCNs innervate different neurons in the striatum (STR) and contribute to exploratory motor behavior (Patel et al., 2012) and action strategies (Dautan et al., 2020). In the ascending ventral circuits, previous studies mainly focus on the cholinergic modulation on the ventral tegmental area (VTA) and substantia nigra (SN), which make great contributions to the reward (Steidl et al., 2017; Zhang et al., 2018), addiction (Picciotto et al., 2002; Schmidt et al., 2009), locomotion (Xiao et al., 2016), depressive-like behaviors (Fernandez et al., 2018) and food taking (Dickson et al., 2010). In the descending circuits, PTCNs govern the activities of the pontine reticular nucleus and contribute to various functions, including inhibiting ongoing movement (Takakusaki et al., 2016) and mediating prepulse inhibition of startle (Azzopardi et al., 2018; Jones et al., 2004). They also modulate breathing via projecting to the retrotrapezoid nucleus (Lima et al., 2019).

There are approximately 2000 PTCNs on one hemisphere of the mouse brain (Li et al., 2018). How can the pontine-tegmental cholinergic system innervate so many brain regions to involve in various functions with such a limited number of neurons? Notably, activating of PPN cholinergic neurons have opposite roles in locomotion in the ascending and descending circuits (Dautan et al., 2016; Takakusaki et al., 2016), but the relation of these two circuits is unclear. Although previous studies have mapped the cholinergic fibers in the thalamus (I. Huerta-Ocampo et al., 2020; Sokhadze et al., 2021), striatum, and substantia nigra (Mena-Segovia & Bolam, 2017) and acquired the whole-brain projections of PPN cholinergic neurons (Henrich Martin et al., 2020), we are still confused about the ascending and descending fibers belong to different groups or same group of neurons, which are urgent to be answered at the single-neuron level. Previous studies (Mena-Segovia et al., 2008) reconstructed partial axons in slices and give preliminary evidence that individual PPN neuron has complex axonal fibers, but we until lack the unabridged single-cell connection of pontine-tegmental cholinergic system due to the limited techniques for tracing and imaging.

Herein, combining the sparse labeling with fluorescence micro-optical sectioning tomography (fMOST) serial technologies (Gong et al., 2016), we acquired the morphological atlas and uncovered the projection logic of PTCNs at the single-cell level.

## Results

### Sparse labeling and single-cell reconstruction

To obtain the fine morphology of individual cholinergic neurons, we employed the Cre-dependent virus for sparse labeling (Sun et al., 2020), fMOST system for whole-brain imaging (Gong et al., 2016), and GTree software for single-cell reconstruction (Zhou et al., 2020) (**Figure 1A, B**). Firstly, we performed 100 nl CSSP-YFP on the PPN or LDT of ChAT-Cre mice. Four weeks later, the infected mice were sacrificed and we checked the object regions. The immunofluorescent staining on the slices verified the cell types of labeled neurons (Figure 1—figure supplement 1 A-C) that more than 96% of them were ChAT-positive (from 3 mice). These results indicated that the virus we used has good specificity.

**Figure 1.**
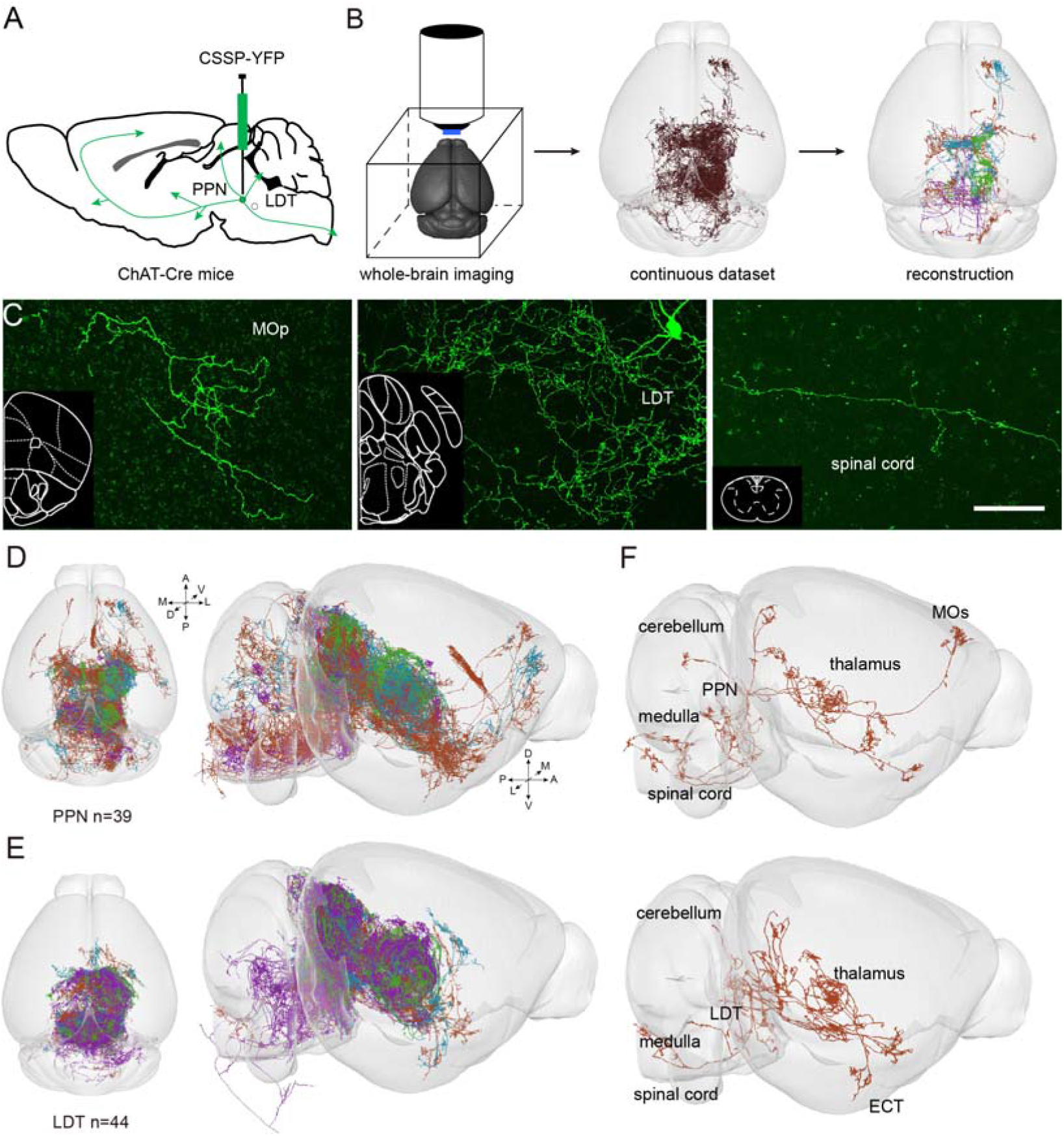
Reconstructions of the individual cholinergic neuron in midbrain. (A) Diagram of the sparse labeling with virus. (B) The main steps for whole-brain data acquisition and single-cell reconstruction. (C) Clear view of Labeled fibers in different regions, such as the cortical area, inject site and spinal cord. (D) (E) Reconstructed neurons mapped to the Allen CCFv3 in 3D. PPN, n = 39 neurons; LDT, n = 44 neurons. Left is plan view of reconstructed neurons. Right is side view of reconstructed neurons in the whole-brain. 83 neurons were reconstructed from six brains. (F) Single neuron stretched its axonal fibers to the cortical areas, thalamus, cerebellum, medulla and spinal cord simultaneously.

Then, we embedded the whole-brain samples with resin and acquired the continuous datasets with high resolution at 0.32 × 0.32 × 1 μm^3^ via fMOST (Gong et al., 2016). To verify the quality of datasets for single-cell reconstruction, we valued the labeled signals in the whole brain. We checked the labeled fibers in different targeting areas (**Figure 1C;** Figure 1—figure supplement 1 D-Q). We found the labeled signal had a good signal-to-noise ratio with the background and single fibers could be distinguished from each other, whether the fibers were in dense or sparse regions. Combining continuous datasets with semi-automatic reconstruction methods (Zhou et al., 2020), we reconstructed 83 supposed cholinergic neurons (PPN, 39 neurons; LDT, 44 neurons) in the pontine-tegmental cholinergic system (**Figure 1D and E**; Figure 1—figure supplement 2). The fibers of reconstructed neurons widely distributed in the whole-brain and some neurons even sent divergent fibers to cortical areas, cerebellum and medulla simultaneously (**Figure 1F**). These fibers covered the main projection areas of PTCNs (Peilin Zhao et al., 2022), which indicated these reconstructed neurons present the morphological characterization of individual cholinergic neurons in the midbrain.

### The whole-brain projection logic of individual PTCNs

To understand the projection patterns of individual PTCNs, we registered all these neurons to the Allen CCFv3 (Wang et al., 2020) and analyzed the terminal properties in the targeting regions (**Figure 2A**). Along with previous studies (Mena-Segovia & Bolam, 2017), we divided the targeting regions into two circuits, that the midbrain and rostral regions into ascending circuit while the pons and medulla into descending circuit. As shown in **Figure 2A**, all reconstructed neurons sent abundant axonal fibers to multiple areas and most of them projected to various nuclei in the ascending and descending circuits simultaneously, except two neurons in the PPN. Moreover, we found most of the reconstructed neurons sent projections to the thalamus, indicating that the thalamus is the major target of both PPN and LDT (PPN, 38/39; LDT, 43/44). Meanwhile, our results showed that many cholinergic neurons projected their axon branches to the cerebellum (PPN, 19/39; LDT, 20/44), and some even extended to the paraflocculus (PFL) (Figure 2—figure supplement 1A).

**Figure 2.**
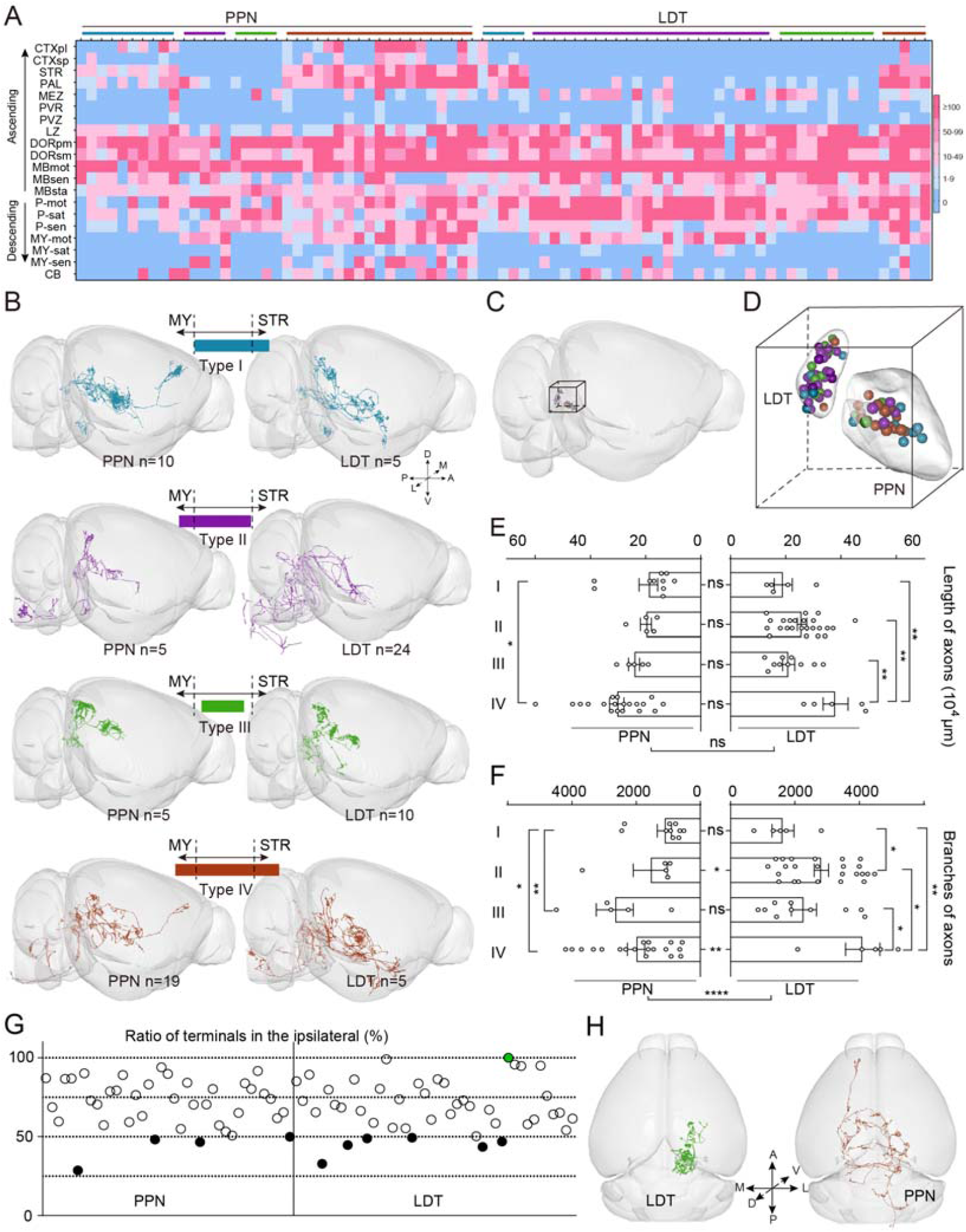
The whole-brain projection patterns of PTINS. (A) The distribution of axonal terminals of single neuron. Each column displayed one reconstructed neuron. Boxes in different color explained the number of terminals of a single neuron in different brain regions. Different color on the top represent reconstructed neurons were classified into different type base on axonal fibers. (B) 3D view of typical neurons with different projection patterns. (C)The soma of all reconstructed neurons in the whole-brain outline. (D)Clear view of reconstructed neurons. The somas in the PPN and LDT neurons were isolated. The soma of neurons with different projection patterns were mixed together. (E) Quantification and comparison of axon length of reconstructed neurons in four types. (F) Quantification and comparison of axon branches of reconstructed neurons in four types. (G) The ratio of axonal terminals in the ipsilateral of single neuron. Green dot showed a neuron confined its axons in the ipsilateral totally. Black filled dots represent 10 neurons had richer axons in the contralateral areas. (H) Left was the top view of ipsilateral restricted neuron. Right was top view of typical contralateral preference neuron. Data shown as Mean ± SEM. two-tailed t-tests, *P < 0.05, **P < 0.01, ***P < 0.001, ****P < 0.0001. The details of abbreviations for brain regions see Supplementary file 1.

We noted that most individual PTCNs extended fibers to the three major certified trajectories (Mena-Segovia & Bolam, 2017) simultaneously, so we cannot analyze the single-cell projection patterns with traditionally defined trajectories. In view of previous studies on the PTCNs mainly focus on the targets between the STR (Dautan et al., 2020) and medulla (Lima et al., 2019) and most reconstructed neurons projected to the interbrain, midbrain and pons concurrently. To furtherly investigate the projection patterns of PTCNs, we classified the reconstructed neurons into different groups and analyzed their characterizations respectively. Depending on whether they target the areas in ascending circuit and descending circuit, such as STR and medulla, four types were distinguished (**Figure 2 A, B**; Supplementary file 2.). As shown in **Figure 2A, B**, type I neurons tended to target anterior telencephalon with axons stretched to the STR but not medulla (PPN, 10 neurons; LDT, 5 neurons). Type II neurons preferred projecting to the posterior brainstem, including the medulla even the spinal cord but not STR (PPN, 5 neurons; LDT, 24 neurons). Neurons of type III projected restrictedly that the axons confined between the STR and medulla (PPN, 5 neurons; LDT, 10 neurons). Type IV neurons had the most widely fibers to both the STR and medulla (PPN, 19 neurons; LDT, 5 neurons). Relatively, the PPN sent richer cholinergic fibers to ascending targets (type I and IV, PPN 29/39; LDT 10/44) while the LDT preferred descending areas (type II and IV PPN 24/39; LDT 29/44). Furthermore, PPN neurons sent more divergent axonal fibers compared with LDT (type IV, PPN, 19/39; LDT, 5/44). Then we investigated the location of somas (**Figure 2 C, D**) and found that the cell bodies of neurons with different projection patterns in the same nucleus were mixed together (Figure 2—figure supplement 1B), which meant adjacent PTCNs might have diverse projections. This agrees with our previous results on the cholinergic neurons in the basal forebrain (Li et al., 2018).

To value the difference of neurons in various projection patterns, we counted the length and branches of reconstructed axons (**Figure 2 E, F**). We found that both PPN and LDT neurons had abundant fibers with a length range from 60 to 90 cm (**Figure 2E**). Then, the length of fibers had no significant difference in total or different types between PPN and LDT. However, in PPN, type I had shorter fibers compared with type IV (PPN, I vs IV, P = 0.0135), while for four types in LDT, type IV had longest axonal fibers (LDT, IV vs I, P = 0.0092; IV vs II, P = 0.0076; IV vs III, P = 0.002). As the distribution of axons and synaptic connection does not always correlate (I. Huerta-Ocampo et al., 2020), we then compared terminal branches of reconstructed neurons. As shown in **Figure 2F**, individual PPN and LDT neurons had a large amount branches ranging from 500 to 5200 and LDT neurons had richer branches in total (PPN vs LDT, P < 0.0001) or in different projection patterns (II, P = 0.029; IV, P = 0.0018). Among different types, type I in the PPN had fewer branches compared with type III and IV (PPN, I vs III, P = 0.0092; I vs IV, P = 0.039) while type IV in the LDT had the richest branches compared with others (LDT, IV vs I, P = 0.0046; IV vs II, P = 0.0.265; IV vs III, P = 0.0177).

In the whole brain perspective, most PTCNs tended to extend fibers in the bilateral hemispheres. To investigate the pattern projecting to bilateral hemispheres, we counted the terminals in the targeting regions of single neurons and quantified the ratio of bilateral axons in the ipsilateral axons. As shown in **Figure 2G**, most neurons projected to the bilateral regions except one neuron in the LDT that confined its axons in the ipsilateral areas. Meanwhile, we also found some neurons (PPN, 4 neurons; LDT, 6 neurons; Figure 1—figure supplement 2) preferred contralateral hemisphere and sent richer fibers to the contralateral (**Figure 2H**).

The dendrites are portals of neurons receiving information from others. We found that different projection patterns of PTCNs in the PPN and LDT had rich dendrites (**Figure 3B**). Consistent with previous studies (Baksa et al., 2019), we divided the dendrites with bipolar and multipolar dendritic trees according to the distribution of longer dendrites (**Figure 3B**) and found a few of them had bipolar dendritic trees while most were multipolar (Figure 3—figure supplement 1). Then we analyzed the spatial distribution of dendritic branches with the Sholl analysis (**Figure 3C**). Our results suggested that the dendrites of both PPN and LDT neurons were mainly distributed in about 600 μm away from the soma and most dendrites gathered in the 50-350 μm and the LDT neurons had richer dendrites. To confirm the result, we quantified the length and branches of all reconstructed dendrites. As shown in **Figure 3D, E**, LDT neurons had significant longer dendrites and more branches in total (P < 0.0001) and different types, especially the type I (PPN vs LDT: dendrites, P = 0.0093; branches, P = 0.0252), type II (PPN vs LDT: dendrites, P = 0.0273; branches, P = 0.0129), and type V (PPN vs LDT: branches, P = 0.004). While in same nuclei, different types of neurons had similar dendrites and branches in quantification, except the type III in the PPN, which had longer dendrites than type I (P = 0.0098) and V (P = 0.0465), and type II in the LDT, which had longer dendrites and richer branches than type III (dendrites, P = 0.0083; branches, P = 0.0231). These results suggested that different types of PTCNs had similar dendrites but the LDT cholinergic neurons had richer dendrites in comparison with PPN.

**Figure 3.**
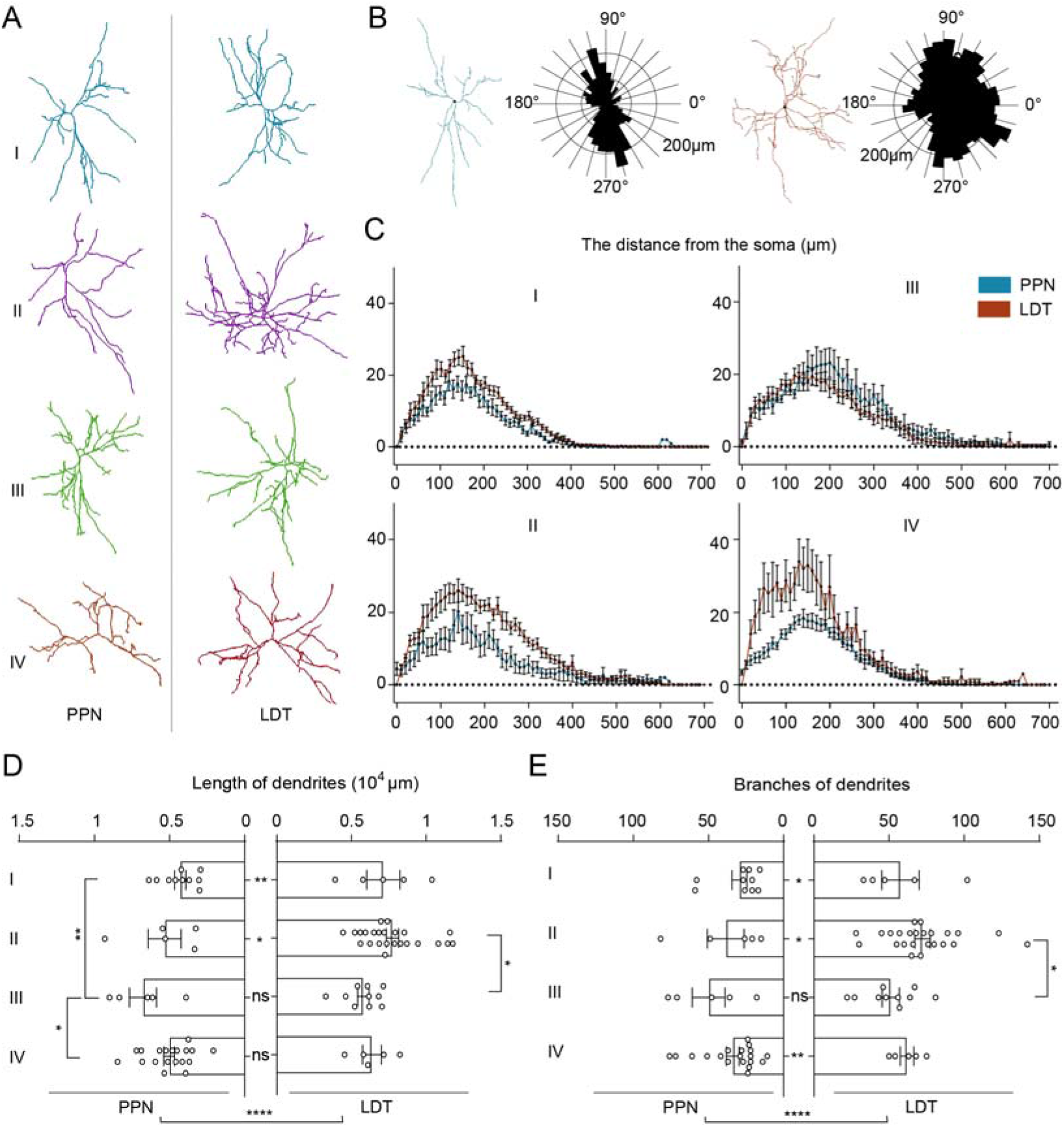
The dendritic morphology of PTCNs. (A) Typical dendrite of PTCNs in different projection patterns. (B) Polar analysis of dendrite. Left was dendritic tree of a cholinergic neuron with bipolar dendritic tree. Right was dendritic tree of a cholinergic neuron with multipolar dendritic tree. (C) Sholl analyses of dendrites of PTCNs in different projection patterns. (D) (E) Quantification and comparison of length and branches of reconstructed dendrites. Data shown as Mean ± SEM. two-tailed t-tests, *P < 0.05, ****P < 0.01, ****P < 0.0001.

In a word, the fancy morphologies rebuilt in the whole brain certify that individual PTCNs send abundant fibers to multiple areas in the bilateral hemispheres with various projection patterns. The PPN neurons project more divergent axon fibers to ascending and descending regions while LDT neurons contain richer axonal branches and dendrites.

### The projection logic of PTCNs in distinct circuits

In the ascending circuits, the thalamus is one the most important pathways of the pontine-tegmental cholinergic system for information dissemination. The PTCNs involve auditory sensation (F. Luo & Yan, 2013), sensorimotor (Muller et al., 2013), and spatial memory(Mitchell et al., 2002) via innervating different thalamic nuclei.

Single-cell morphology suggested that most PTCNs sent abundant fibers to the thalamus (**Figure 2A**). To explore the cholinergic projection logic in the thalamus, we divided the bilateral thalamus into 22 subregions according to the Allen CCFv3 and then counted the terminals of individual neurons (Figure 4—figure supplement 1A; Supplementary file 3). Consistent with output atlas, both PPN and LDT cholinergic neurons extended rich fibers in the bilateral thalamus, especially the anterior, ventral, and medial areas. Meanwhile, similar to the synaptic distribution in rats (I. Huerta-Ocampo et al., 2020), individual LDT cholinergic neurons preferred the thalamic limbic nuclei, such as the anterior group of the dorsal thalamus (ATN). The single PTCNs tended to innervate multiple thalamic nuclei that more than 80% of reconstructed neurons projected to at least five thalamic nuclei simultaneously (**Figure 4A**). As some individual PTCNs sent axonal fibers to the bilateral thalamus, to verify the organization of cholinergic fibers in the bilateral thalamus, we quantified the proportion of axonal terminals in the ipsilateral thalamus (Figure 4—figure supplement 1B). In the thalamus-projecting PTCNs, we found that about a quarter of them only innervated ipsilateral thalamic nuclei (**Figure 4B**; Figure 4—figure supplement 1B) while others targeted the bilateral thalamus synchronously. Meanwhile, we also found about a tenth of PTCNs preferred the contralateral thalamus (**Figure 4C, D**). These results suggested that there had three distinct patterns of dominance from the PTCNs to the bilateral thalamus (**Figure 4D**).

**Figure 4.**
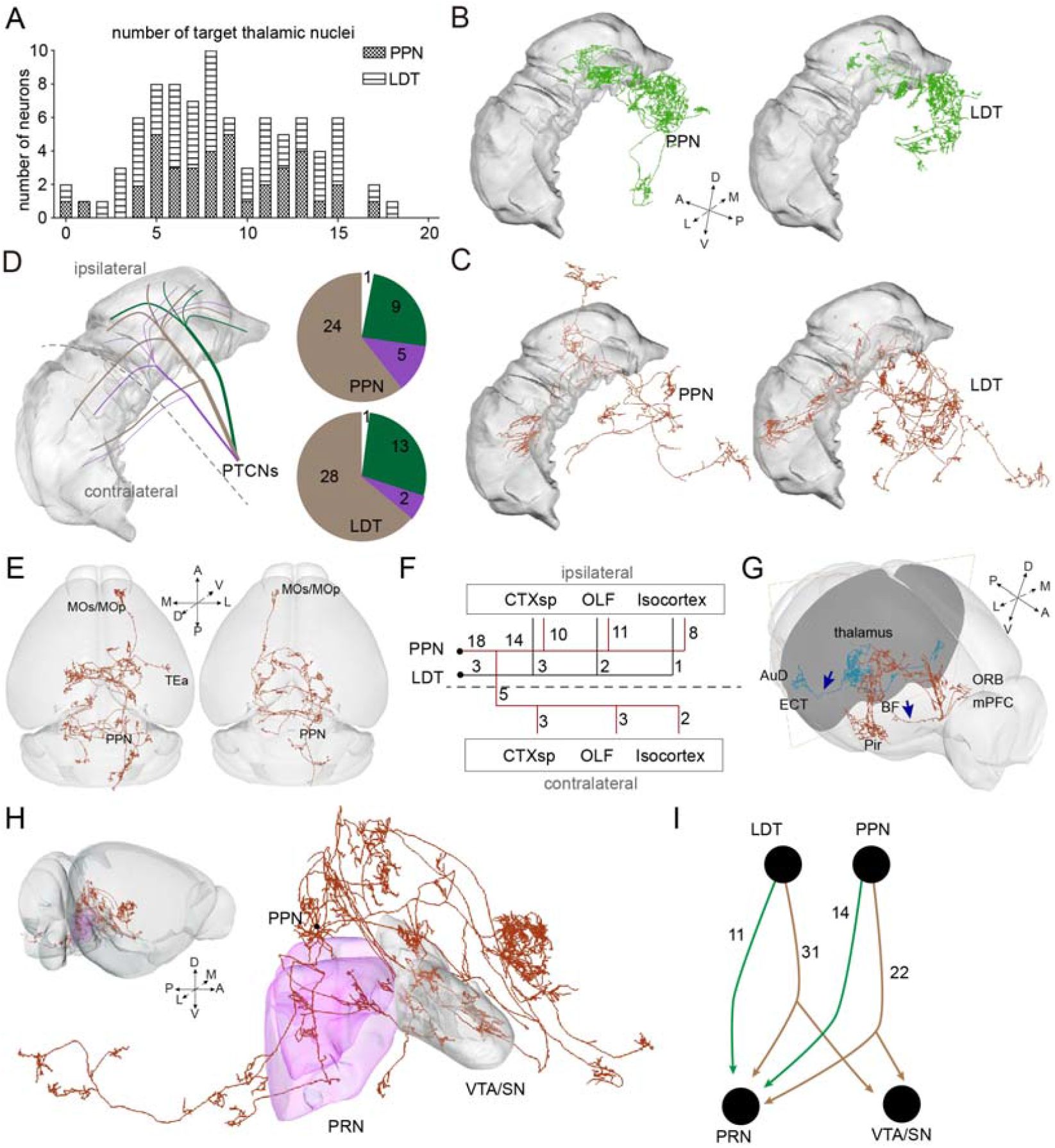
The projection logic in ascending and descending circuits. (A) Number of thalamic nuclei innervated by individual PTCN. (B) Typical neuron only innervating the ipsilateral thalamus. (C). Typical neuron preferred the contralateral thalamus. (D) Schematic diagram and number of three patterns of PTCNs innervating the thalamus. (E) 3D view of typical neurons projecting to the ipsilateral (left) and contralateral (right) cortical areas. (F) The targeting areas of cortical-projecting neurons. (G) Two separated pathways projecting to rostral and lateral cortical areas. (H) 3D view of typical neuron projecting to the PRN and VTA/SN simultaneously. (I) Number of reconstructed PTCNs projecting to the PRN and VTA/SN. The details of abbreviations for brain regions see Supplementary file 1.

The cortical areas are the advanced center of the nervous system and the thalamus is the main source of subcortical information in the cortical regions (Oh et al., 2014; P. Zhao et al., 2020). Previous studies have certified that the activities of PTCNs influence the function of cortical neurons via indirect circuits (F. Luo et al., 2011; Valencia et al., 2013), but we know little about the direct pathways. Our results revealed stably projection from PTCNs to different cortical areas in the bilateral hemispheres (**Figure 4E**) and most of them originated from the PPN (**Figure 4F**).

Then we analyzed the cortical projection neurons and found there were two separate pathways extending to cortical areas. Some axons stretched through the basal forebrain and projected to cortical areas in the anterior, while other fibers came through the lateral thalamus and reached the lateral cortex, such as the perirhinal areas (**Figure 4G**). Meanwhile, we found the PTCNs targeting the cortex also co-projecting to thalamic nuclei (**Figure 2A**).

In the descending circuits, the fibers of PTCNs were widely distributed in the pons and medulla, and some axons extended to the spinal cord. Previous studies have certified that activating of PPN cholinergic neurons in ascending circuits and descending circuits make opposite impacts in locomotion (Dautan et al., 2016; Takakusaki et al., 2016). We analyzed the relation of these two functionally different circuits based on the single-cell morphology atlas. As shown in **Figure 4H, I**, most of the reconstructed PPN cholinergic neurons (36/39) targeted the PRN, and more than half them (22/39) co-projected to the VTA/SN. Here was a surprise that we did not find a single neuron innervating the VTA/SN alone without co-projecting to PRN. Then, we also investigated the cholinergic circuits from the LDT to VTA/SN and PRN. Similar to the PPN, the majority of LDT neurons (42/44) extended fibers to the PRN, and about three-quarters of them co-projected to the VTA/SN. These results meant that the cholinergic modulation from the PTCNs to the VTA/SN often together with co-projecting to the PRN and there existed another group of PTCNs innervated the activity of PRN alone.

## Discussion

The PTCNs send abundant fibers to multiple regions in the ascending and descending circuits and participate in various functions. We presented here the detailed morphology of PTCNs in the whole brain and uncovered the projection logic at single-cell level. Individual PTCNs always projected to ascending and descending circuits simultaneously and could be divided into four groups with different projection patterns. In the ascending circuits, individual PTCNs innervated the bilateral thalamus with three distinct patterns according to the axonal selection in the bilateral thalamus. Moreover, PTCNs projected directly to the cortical areas with two separate pathways.

### Surprising complex morphology of PTCNs

Why do we need the morphology of single-cell? The output atlas indicates that projecting neurons involve in diverse circuits and functions via complex axonal connections with various nuclei, but we are still confused about how these neurons are organized together, such as whether the neurons innervating different regions belong to the same or separate groups. The morphological studies on individual neurons indicated that a single projecting neuron always has abundant fibers in multiple targeting areas with projection preferences. For example, the pyramidal neurons in the cortex can be subdivided into three types (intratelencephalic, pyramidal, and contralateral tract) with distinct projection targets and functions (Peng et al., 2021). Individual cholinergic neurons in the basal forebrain also have abundant fibers and could be gathered into different types due to diverse projection preferences to the olfactory bulb, cortical areas, and hippocampus (Li et al., 2018). Similarly, with the projection logic, we gathered the individual PTCNs into four types with different lengths and extended ranges of axons. Morphological results in the 3D showed that single PTCNs contained rich fibers and simultaneously innervated tens of nuclei ranging from cortical areas to the cerebellum and medulla with the axonal length at 9 ∼ 60 cm and 500 ∼ 5000 terminal branches. In particular, it was quite striking that almost every neuron projected to multiple nuclei in the midbrain, thalamus, and pons. We have even found several neurons deliver their cholinergic fibers to the cortex, thalamus cerebellum, and spinal cord simultaneously. Our results indicated that PTCNs projected to the three major axonal trajectories (Mena-Segovia & Bolam, 2017) mainly from the same groups rather than segregate clusters. To some extent, this may explain how the pontine-tegmental cholinergic system participates in a variety of physiological functions with a finite number of cholinergic neurons (Li et al., 2018).

In summary, we may investigate the complexity of PTCNs mainly in the following two aspects. On the one hand, most reconstructed neurons targeted ascending and descending areas synchronously, which was not consistent with previous views (Mena-Segovia & Bolam, 2017) that thought only a small ratio of PPN neurons targeting descending areas and mainly were non-cholinergic. Actually, from the reconstruction of individual neurons, we found that although the PTCNs projected to the medulla with relatively fewer fibers (Henrich Martin et al., 2020), more than half of them sent fibers to the medulla. On the other hand, we also found some neurons sent richer fibers to the contralateral regions, which might deliver information to the bilateral hemisphere. Axonal selection of ipsilateral and/or contralateral targets is crucial for integrating bilateral information and coordinated movement. The PTCNs decrease significantly in PD patients (Sebille et al., 2019), who always are abnormalities of movement, including tremors, difficulties with gait and balance (Mazzoni et al., 2012). The PPN is an important clinical target for deep brain stimulation in PD patients (Nowacki et al., 2019), which reminds us that PTCNs may play an important role in somatic homeostatic regulation via projecting abundant fibers to bilateral hemispheres.

### Unique ascending and descending projection patterns of PTCNs

The ascending and descending information flow is highly ordered in the central nervous system. As an important interaction area for ascending and descending information transmission, some midbrain neurons preferentially dominate the ascending or descending circuits (Beier et al., 2015; Ferreira-Pinto et al., 2021; Xu et al., 2021). Cholinergic neurons in the midbrain send abundant fibers to various regions in the ascending and descending circuits. In anticipation, we thought that these cholinergic neurons might be grouped into different groups for ascending and descending projections, like the glutaminergic neurons in the mesencephalic locomotor region (Ferreira-Pinto et al., 2021). Surprisingly, the morphology of individual PTCNs showed an unparalleled complexity that most of them sent abundant fibers to ascending and descending areas simultaneously. This means that when PTCNs send information to ascending regions, they often innervate descending areas via collaterals.

Moreover, previous studies (Dautan et al., 2016; Takakusaki et al., 2016) have certified that activating of PPN cholinergic neurons play opposite roles in locomotion via ascending and descending circuits, which mean these two circuits may belong to different groups of cholinergic neurons. However, our results indicated that all the PTCNs projecting to the VTA/SN had co-projection fibers to the PRN while some PTCNs innervated the PRN alone without fibers to the TVA/SN. All of which showed that the opposite functions of two cholinergic circuits might come from different neurons, or from the different responses of downstream neurons to the same neurons. These results hint to us that PTCNs modulating many ascending and descending circuits may mainly originate from the same group of neurons. However, it also raises a larger question, as a region for information interaction, why PTCNs dominate the ascending and descending circuits simultaneously and what it means in the flows of information in the midbrain. More experiments are needed to decode the functional difference of ascending and descending collaterals from the same neurons.

### Projection logic of cholinergic fibers ascending to the thalamus and cortex

The thalamus is an important relay station for sensorimotor information in mammals, and act as the main pathway for the pontine-tegmental cholinergic system delivering information in ascending circuits. As the main source of acetylcholine for the thalamus (M. M. Mesulam et al., 1989; Motts et al., 2010; Paré et al., 1988; Sofroniew et al., 1985a), PTCNs send fibers to different thalamic nuclei and form abundant synaptic connections (I. Huerta-Ocampo et al., 2020; Sokhadze et al., 2021).

In our studies, we uncovered the projection logic from the PTCNs to the thalamus and found the thalamus-projecting circuits mainly presented the following characteristics. For the first, most PTCNs innervated the thalamic nuclei directly. Only two of 83 reconstructed neurons did not project to the thalamus, the ratio was higher than that retrograde traced from the thalamus (Sofroniew et al., 1985b). These results are not contradictory due to that retrograde tracing had trouble infecting the cholinergic neurons with sparse fibers in specific thalamic nuclei. For the second, individual PTCNs tended to govern multiple thalamic nuclei synchronously. We found more than 80% of reconstructed PTCNs innervated at least five thalamic nuclei and it was amazing that some of them even projected to 18 nuclei in the bilateral thalamus simultaneously. This means that the regulation of PTCNs to the thalamus is mainly diffuse rather than concentrated in a few regions. As we know, PTCNs modulate different thalamic nuclei (F. Luo & Yan, 2013; Mitchell et al., 2002; Muller et al., 2013) which integrated various sensorimotor information and delivered it to diverse cortical areas (Oh et al., 2014; P. Zhao et al., 2020). We speculate that individual PTCNs always send complex information to various thalamic nuclei, which sort the information and then deliver it to the specific cortex. Finally, we decoded three projection patterns from the PTCNs with different preferences in the ipsilateral and/or contralateral thalamus. Hinting that there may be three different modes of information transmission from the pontine-tegmental cholinergic system to the thalamus for further information progress. Specifically, a large amount of PTCNs dominated the activity of the bilateral thalamus and some of them even preferred the contralateral. They may play an important role in coordinated movement and contribute to some diseases with movement disorders, such as PD (Pepeu et al., 2017) and epilepsy (Soares et al., 2018), the cholinergic circuits in the thalamus of these patients present abnormal activity (Miller et al., 1991; Muller et al., 2013).

The cortical areas are the higher nervous center of mammals. Various studies have indicated that PTCNs govern the activity of cortical neurons via indirect pathways such as the thalamus (F. Luo et al., 2011). In this study, we certified the direct innervation from the PTCNs to the bilateral cortex and there are two separate pathways to the anterior and lateral cortical areas. The cortex-projection PTCNs also presented stable co-projection to various thalamic nuclei, which meant that direct regulation of the cortex by PTCNs was often accompanied by indirect pathways through the thalamus. Although only a small portion of the PTCNs innervated the activity of cortical neurons directly, it was important for understanding the information transmission of the pontine-tegmental cholinergic system in ascending circuits.

### Morphological difference of PPN and LDT cholinergic neurons

As two components of the pontine-tegmental cholinergic system, PPN and LDT cholinergic neurons have their characteristics in connections and functions (Dautan et al., 2016; Icnelia Huerta-Ocampo et al., 2021; Xiao et al., 2016). Previous studies have certified that the PPN and LDT cholinergic neurons innervate similar regions with different preferences (I. Huerta-Ocampo et al., 2020; Mena-Segovia & Bolam, 2017; Sokhadze et al., 2021). In this study, we certified that the PPN and LDT cholinergic neurons had similar projection patterns at the single-cell level and could be divided into four groups. Comparatively, the cholinergic fibers of PPN neurons presented more divergent to the telencephalon, interbrain, midbrain, and lower brainstem simultaneously, while LDT neurons had richer branches that mainly concentrated in the poster brainstem. In addition, the dendrites of LDT cholinergic neurons had richer branches and longer dendrites than that of PPN. As portals of neurons receiving information, the dendrites influence the number and identity of presynaptic inputs (Lefebvre et al., 2015), which implies that LDT neurons may receive wider afferent information.

In conclusion, we acquired the single-cell morphology of PTCNs on a brain-wide scale and decoded the projection logic in the whole-brain and some specific circuits. We revealed the individual PTCNs sent abundant fibers in multiple nuclei in the ascending and descending circuits synchronously and could be gathered into four types with different projection patterns. We decoded three different projection patterns from PTCNs to the bilateral thalamus and two separated pathways to cortical areas. This study maps the finest morphological atlas of the pontine-tegmental cholinergic system and gives us a better understanding of projection logic of them.

## Materials and Methods

### Key resources table

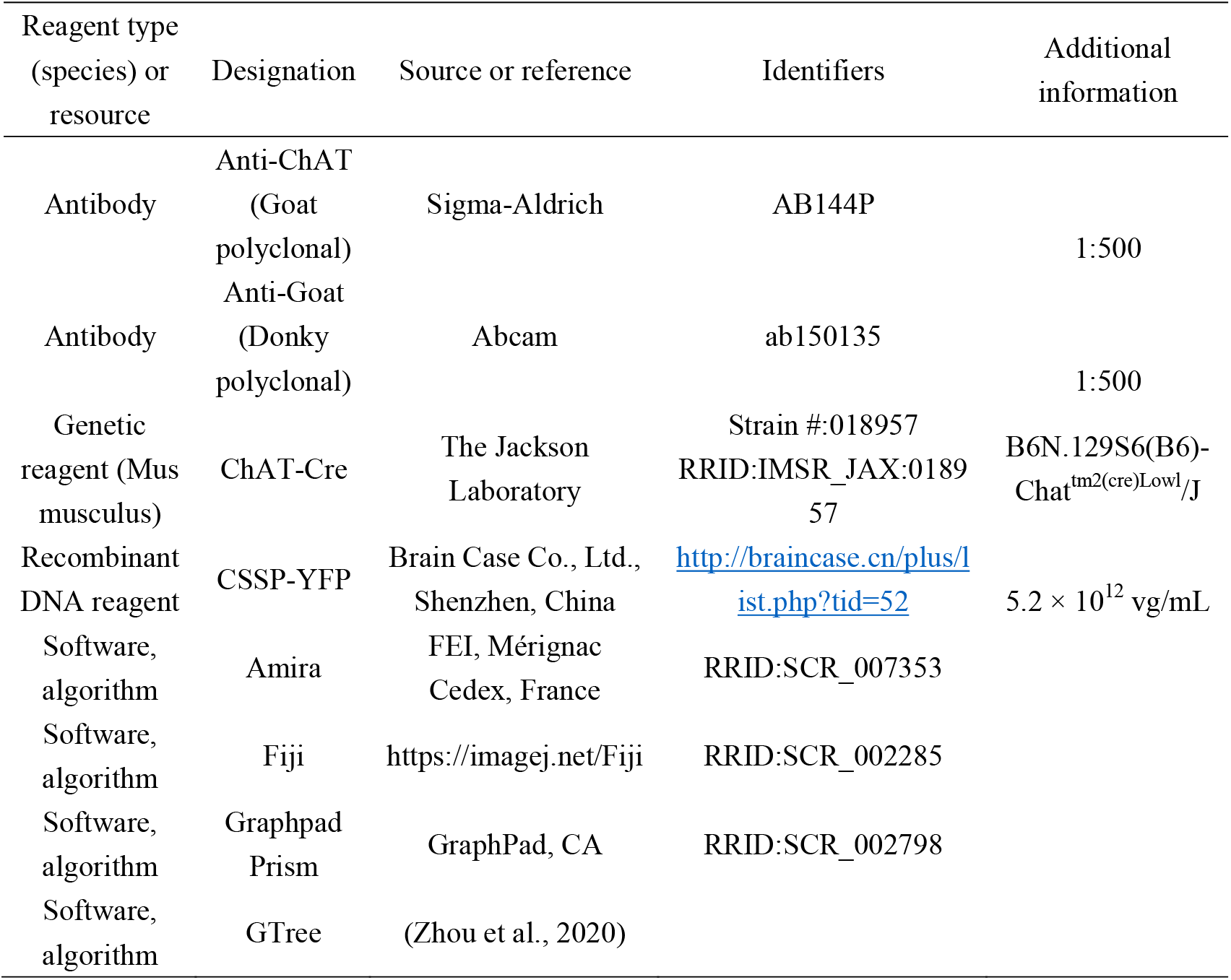

### Animals

Adult (2-4 months) ChAT-ires-Cre mice were used in this study. ChAT-Cre transgenic mice (stock No: 018957) were purchased from Jackson Laboratory. The mice were kept under a condition of 12-hour light/dark cycle with food and water ad libitum. All animal experiments were approved by the Animal Care and Use Committee of Huazhong University of Science and Technology.

### Tracer information

For sparse labeling, we employed the CSSP-YFP (5.2 × 10^12^ vg/mL) packed by Brain Case (Brain Case Co., Ltd., Shenzhen, China). CSSP-YFP virus is produced by co-packaging the rAAV-EF1α-DIO-Flp plasmid and rAAV-FDIO-EYFP plasmid with the ratio of 1:20,000 in single rAAV production step (Sun et al., 2020).

### Surgery and Viral Injection

Before virus injection, we anesthetized the mice with mixed anesthetics (2% chloral hydrate and 10% ethyl urethane dissolved in 0.9% NaCl saline) according to their weight (0.1 ml/10 g). The brain of anesthetized mice was fixed with a stereotaxic holder to adjust the position of skulls. Then a cranial drill (∼ 0.5 mm diameter) was employed to uncover the skulls above the target areas. For sparse labeling, we injected 100 nl CSSP-YFP into the PPN or LDT.

### Histology and Immunostaining

The histological operations followed previous studies (P. Zhao et al., 2020). Shortly, four weeks after AAV injection, anesthetized mice were perfused with 0.01 M PBS (Sigma–Aldrich, United States), followed with 2.5% sucrose and 4% paraformaldehyde (PFA, Sigma–Aldrich, United States) in 0.01 M PBS. Then, the brains were removed and post-fixed in 4% PFA solution overnight.

For immunofluorescent staining, some samples were sectioned in 50 μm coronal slices with the vibrating slicer (Leica 1200S). All sections containing PPN or LDT were selected to characterize the labeled neurons in inject site. These sections were blocked with 0.01 M PBS containing 5% (wt/vol) bovine serum albumin (BSA) and 0.3% Triton X-100 for 1 h at 37 °C. Then the sections were incubated with the primary antibodies (12h at 4 °C): anti-ChAT (1:500, goat, Sigma-Aldrich, AB144P). Then the sections were washed in PBS five times at room temperature. Next, these sections were incubated with the fluorophore-conjugated secondary antibody (1:500, Abcam: Alexa-Fluor 647, donkey anti-goat) for 2 h at room temperature. After rinsing with PBS, DAPI (1 ng/mL) was performed on stained sections for 5 min, and sections were finally mounted after washing. We acquired the stained information of sections with the confocal microscope (LSM 710, Zeiss, Jena, Germany).

### Imaging and 3D visualization

For whole-brain imaging, virus-labeled samples were dehydrated with alcohol and embedded with resin (Ren et al., 2018). Then the whole brain datasets were acquired with the fMOST system. In short, we fixed the sample on the base and acquired the image of the top surface with two fluorescent channels; the imaged tissue was subsequently removed. Thus, we obtained the continuous whole-brain dataset layer by layer with high resolution (0.32 × 0.32 × 1 μm^3^).

For 3D visualization and statistical analysis of whole-brain datasets, we registered the whole-brain datasets to the Allen Mouse Brain Common Coordinate Framework version 3 (Allen CCFv3)(Wang et al., 2020). The methods of registration have been described previously(Ni et al., 2020). Briefly, we employed image preprocessing to correct uneven illumination and then remove background noise. The down sampling data (the voxel resolution of 10 × 10 × 10 μm^3^) was uploaded into Amira software (v6.1.1, FEI, Mérignac Cedex, France) to distinguish and extract regional features of anatomical invariants, including the outline of the brain, the ventricles, and the corpus striatum, etc. Next, the grey-level-based registration algorithm (SyN) was employed to register the extracted features. Basic operations including extraction of areas of interest, resampling, and maximum projection performed via Amira software and Fiji (NIH).

### Morphological reconstruction of single neuron

83 neurons were reconstructed from six brains. For single-cell morphological analysis, we reconstructed the morphology of sparsely labeled neurons with semi-automatic methods followed previous studies (Zhou et al., 2020). Briefly, we acquired the spatial coordinates of labeled somas in high-resolution data and transformed the data format of GFP-labeled data from TIFF to TDI type via Amira firstly.

Then the data block containing the given soma was loaded into GTree software and we assigned the soma as the initial point and marked all its fibers with unfinished tags. Next, we selected one uncompleted fiber and traced it in the next block with automatic tracing. Then we checked the traced fiber and marked its branches with unfinished tags. We repeated the above procedure until the selected fiber was finished and then we reconstructed the remaining unfinished fibers until all the fibers were achieved. The reconstructed neurons were checked back-to-back by three persons.

The tracing results were saved in SWC format. Meanwhile, we registered the PI-labeled data and the corresponding tracing results to the reference atlas with the methods mentioned above. Considering the distribution of axons and synaptic connection does not always correlate(I. Huerta-Ocampo et al., 2020), we counted the terminal branches of reconstructed neurons to represent the connection between single cholinergic neurons and targeting areas.

### Statistical information

For terminals quantification, the reconstructed neurons carried spatial information of all nodes that we could calculate the ended nodes based on registered single neurons to obtain the terminals in different regions. In addition, we employed the Amira software to quantify the length of dendrites and axons of individual neurons. Statistical graphs were generated using GraphPad Prism v.8.02 and Microsoft Excel (Office 2020). We employed GraphPad Prism v. 8.02 for significance test, Neurolucida360 software for polar histogram, and MATLAB (2017a) for the Sholl Analysis. We conducted two-tailed t-tests to compare the difference. The confidence level was set to 0.05 (P value) and all results were presented as mean ± SEM.

## Acknowledgments

This work was supported by the National Science and Technology Innovation 2030 Grant (No.2021ZD0201001), NSFC projects (Nos. 61890953, 32192412, 31871088) and CAMS Innovation Fund for Medical Sciences (2019-I2M-5-014) and the Director Fund of WNLO. We thank Yang Yang and Mengting Zhao from Huazhong University of Science and Technology for help with experiments and data analysis. We thank the Optical Bioimaging Core Facility of HUST for support with data acquisition.

## Author Contributions

Hui Gong, Xiangning.Li and Qingming Luo conceived and designed the study. Peilin Zhao performed the experiments and analyzed the data. Tao Jiang and Jing yuan acquired the continuous whole brain datasets. Huading Wang, Xueyan Jia and Anan Li processed the whole-brain data and reconstructed the neurons. Xiangning Li, Hui Gong, and Peilin Zhao wrote the manuscript.

## Data availability

The analysis results and data have been uploaded in Supplementary Files 1, 2 and 3. The TB-sized raw data of sparse-labeling samples and 3D data of reconstructed neurons can be accessed at http://atlas.brainsmatics.org/a/zhao2206.

## Supplemental Figures

**Figure 1—figure supplement 1.**
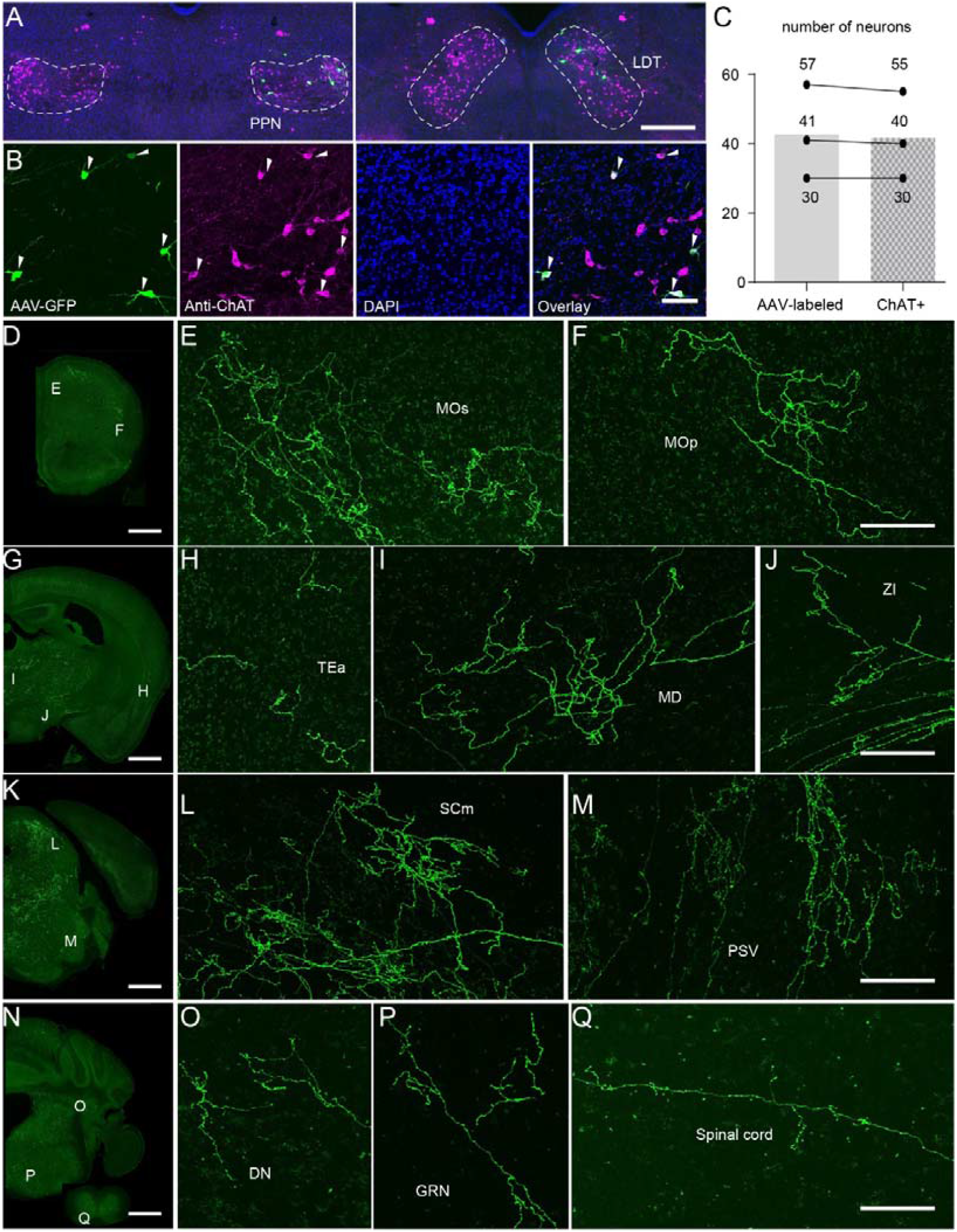
Sparse-labeling of cholinergic neurons. (A) Sparse labeled neurons in the PPN and LDT. (B) A four-panel presentation with AAV-YFP, anti-ChAT, DAPI and merged, arrows point out cholinergic positive neurons. (C) Calculation of all labeled neurons and ChAT+ labeled neurons (n = 3 mice). (D-Q) Labeled fibers ranging from the cortical areas to spinal cord. Scale bar, (A) 500μm; (B) 50μm; (D, G, K, N) 1000μm; (E, F, H, I, J, L, M, O, P, Q) 100μm.

**Figure 1—figure supplement 2.**
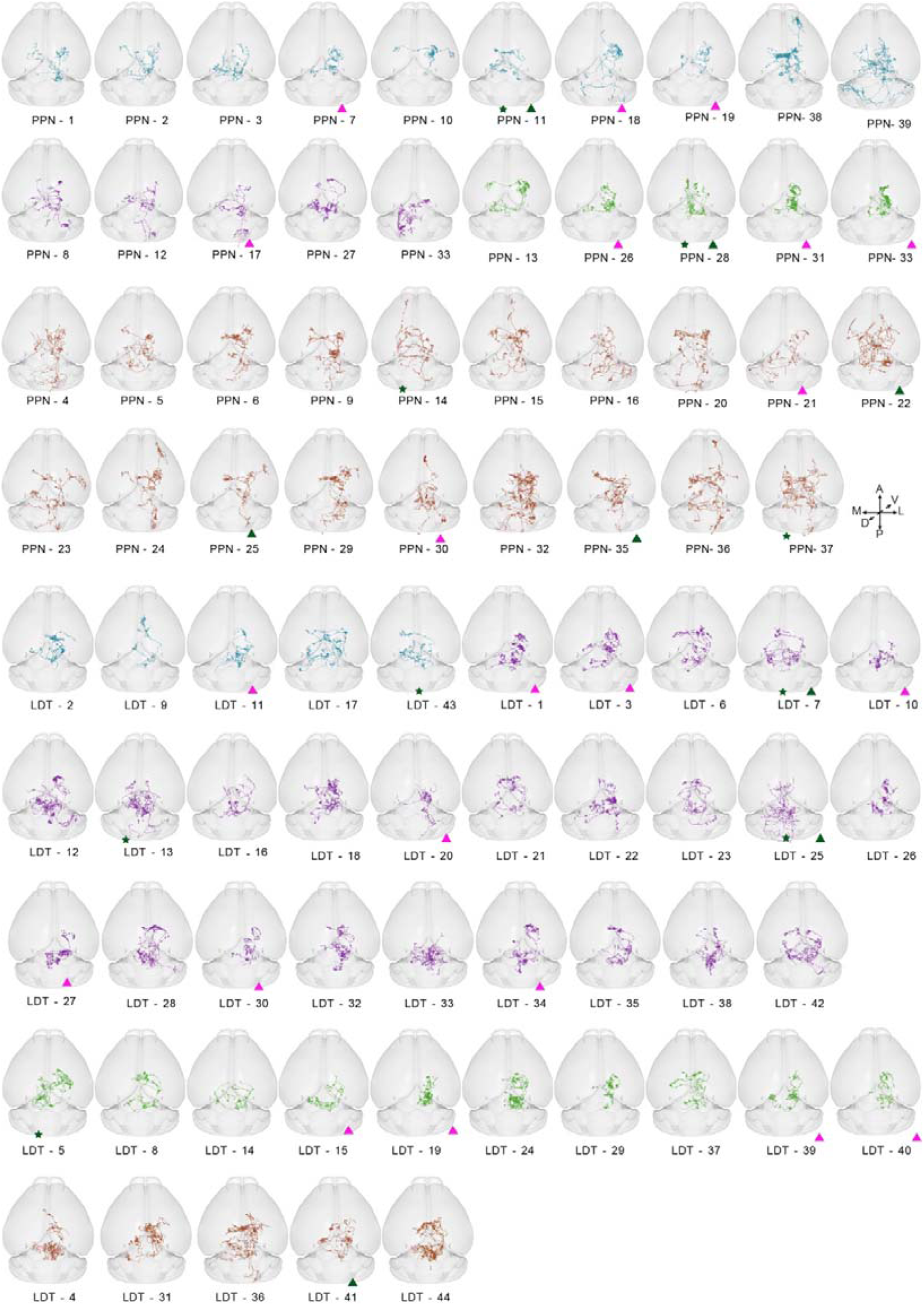
3D view of all 83 reconstructed cholinergic neurons. Pentagons presented neuron had richer axon branches in the contralateral hemisphere. Red triangle pointed neuron restricted its fibers in the ipsilateral thalamus. Green triangle pointed neuron sent richer fibers to the contralateral thalamus.

**Figure 2—figure supplement 1.**
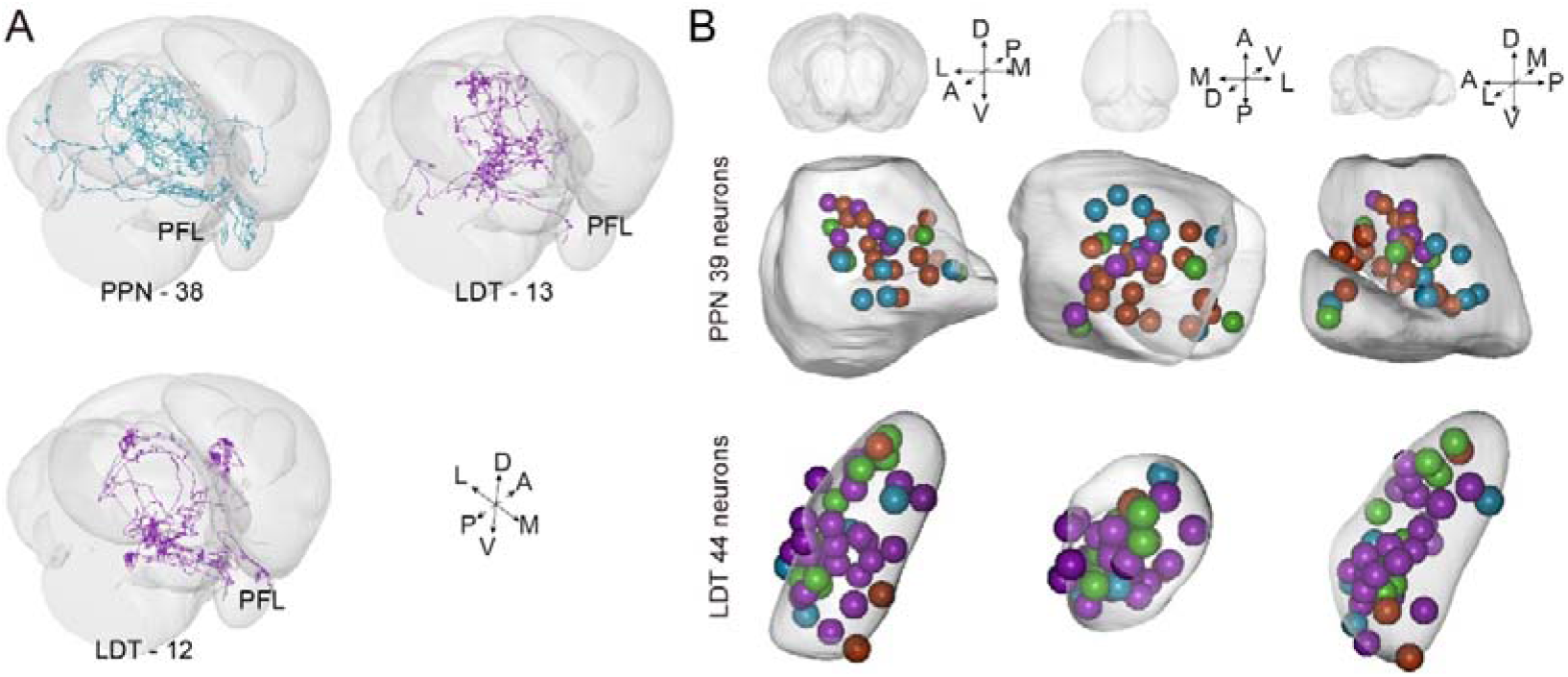
PFL-projection neurons and the soma of reconstructed neurons. (A) 3D view of three neurons projecting to the PFL. (B) 3D view of all reconstructed soma.

**Figure 3—figure supplement 1.**
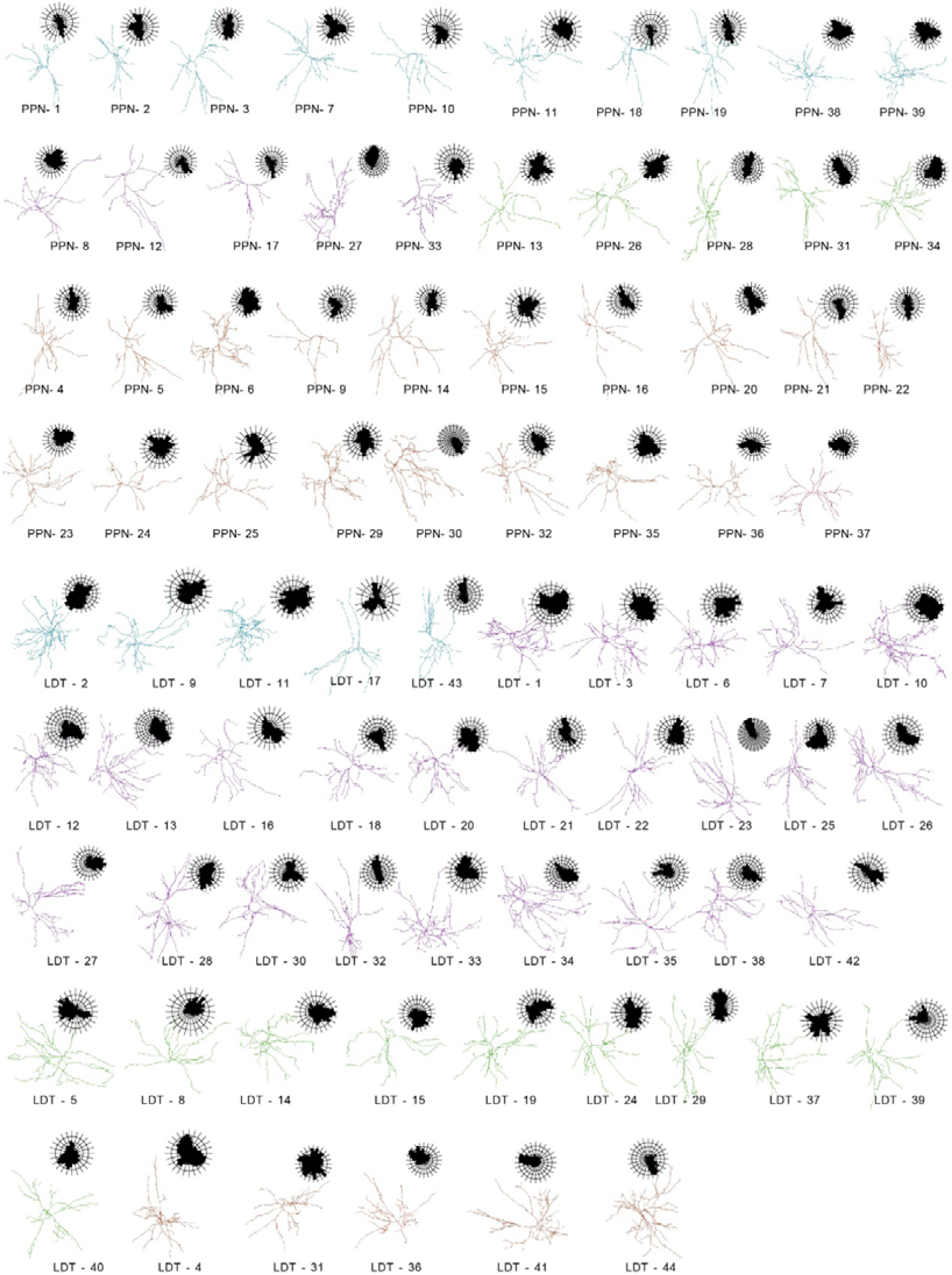
The morphology and polar analysis of 83 reconstructed neurons.

**Figure 4—figure supplement 1.**
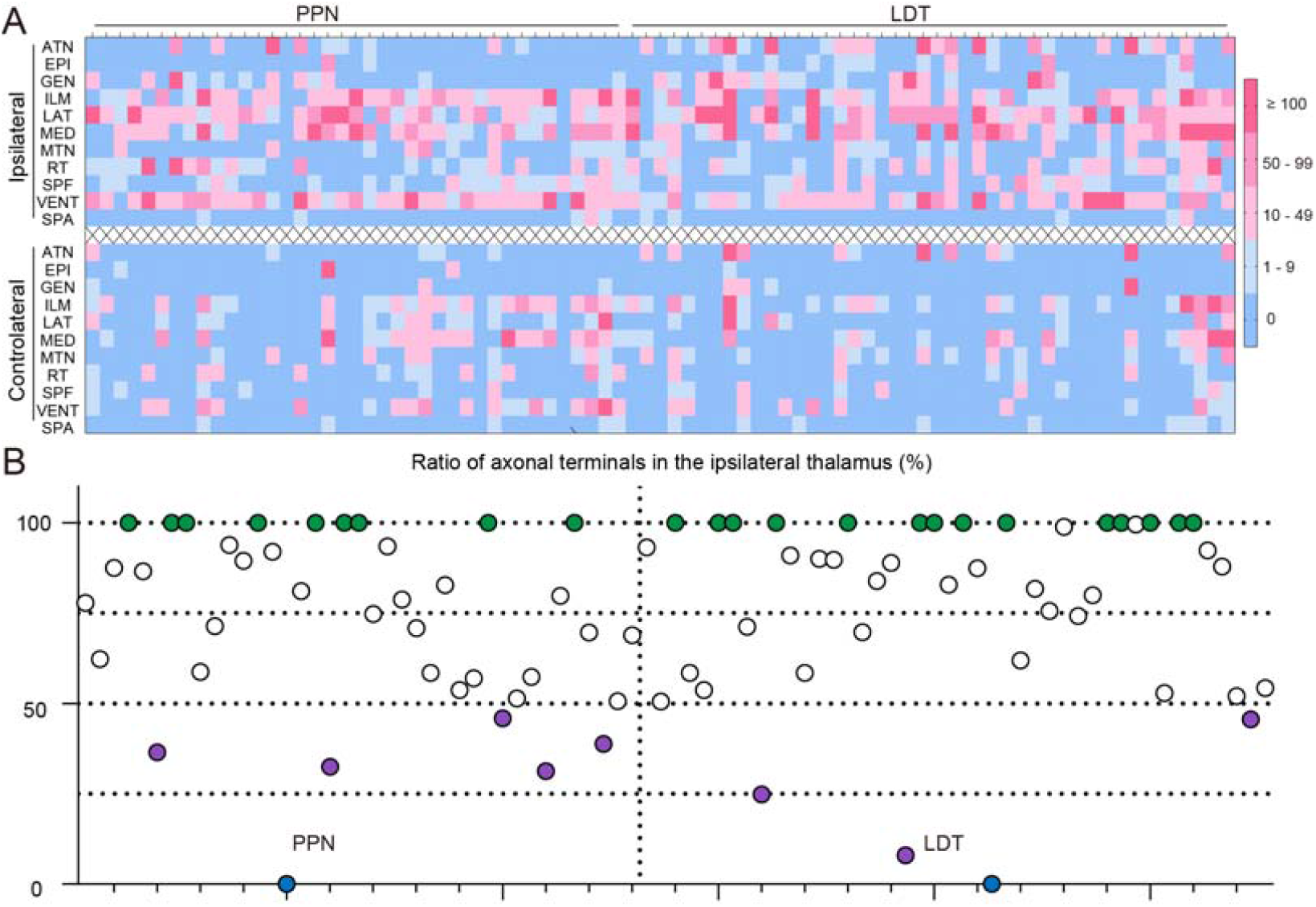
Quantitative analysis of axons of thalamic projection PTCNs. (A) The distribution of axonal terminals in thalamus of single neuron. Each column displayed one neuron. Boxes in different color explained the number of terminals of a single neuron in different brain regions. (B) The ratio of terminals in the ipsilateral thalamus of single neuron. Green dots showed neurons confined its axons in the ipsilateral thalamus. Purple filled dots represent neurons had richer axons in the contralateral thalamus. Blue dots represent neurons did not target the thalamus. The details of abbreviations for brain regions see Supplementary file 1.

